# Fast DNA-PAINT imaging using a deep neural network

**DOI:** 10.1101/2021.11.20.469366

**Authors:** Kaarjel K. Narayanasamy, Johanna V. Rahm, Siddharth Tourani, Mike Heilemann

## Abstract

DNA points accumulation for imaging in nanoscale topography (DNA-PAINT) is a super-resolution technique with relatively easy-to-implement multi-target imaging. However, image acquisition is slow as sufficient statistical data has to be generated from spatio-temporally isolated single emitters. Here, we trained the neural network (NN) DeepSTORM to predict fluorophore positions from high emitter density DNA-PAINT data. This achieves image acquisition in one minute. We demonstrate multi-color super-resolution imaging of structure-conserved semi-thin neuronal tissue and imaging of large samples. This improvement can be integrated into any single-molecule microscope and enables fast single-molecule super-resolution microscopy.

## Introduction

The advent of super-resolution imaging has overcome the diffraction-limited barrier of light microscopy into obtaining images at nanometer spatial resolution. One powerful super-resolution technique for imaging cellular samples is single-molecule localization microscopy (SMLM) which builds on the spatio-temporal isolation of single fluorophores and the precise determination of their position, leading to the reconstruction of a super-resolved image (Sauer and Heilemann, 2017). Methods such as (fluorescence) photoactivated localization microscopy ((F)PALM) (Betzig *et al*., 2006; Hess, Girirajan and Mason, 2006) and (direct) stochastic optical reconstruction microscopy ((*d*)STORM) (Rust, Bates and Zhuang, 2006; Heilemann *et al*., 2008) use photoswitchable fluorophores to obtain a temporally and spatially separated fluorescence signal. Points accumulation for imaging in nanoscale topography (PAINT) (Sharonov and Hochstrasser, 2006) and DNA-PAINT (Jungmann *et al*., 2010) employ transiently binding, low-affinity fluorophore labels for this purpose. Both concepts generate a super-resolved image through the localization of a large number of single emitter positions and achieve a spatial resolution in the range of tens of nanometers.

The tradeoff to acquiring super-resolved images with SMLM is the long image acquisition time. The requirements for an SMLM experiment are sparse and isolated emitters per image and a sufficiently high number of emitters detected over time to reconstruct a cellular structure. These two criteria require a large amount of data generation, hence the long imaging time. Several SMLM studies are focusing on overcoming this limitation using improved localization software (Sage *et al*., 2019), high performance computing and algorithms (Wang *et al*., 2017; Munro *et al*., 2019), or modulating the hybridization times of DNA oligonucleotides (Schueder *et al*., 2019; Civitci *et al*., 2020).

In recent years, various deep learning (DL) tools have emerged to facilitate faster image acquisition in SMLM. The ANNA-PALM neural network predicts a complete super-resolved image from a small set of input frames with incomplete structural features (Ouyang *et al*., 2018). Other neural networks aim to predict 2D and 3D structures from high-density SMLM raw images such as Deep-ULM (van Sloun *et al*., 2021), DECODE (Speiser *et al*., 2021), DRL-STORM (Yao *et al*., 2020), DeepLoco (Boyd *et al*., 2018), and LSPARCOM (Dardikman-Yoffe and Eldar, 2020). DeepSTORM (Nehme *et al*., 2018, 2020) is one such convolutional NN that can be trained to predict single-emitter positions from high-density data to obtain super-resolution images from shorter SMLM movies. The ease-of-use of DeepSTORM was bolstered with its implementation into the ZeroCostDL4Mic platform (von Chamier *et al*., 2021).

DeepSTORM performance is largely dependent on an optimal range of emitter densities. While *(d)*STORM and PALM methods were initially used for DeepSTORM, the exponential decrease in emitter density over acquisition time due to photobleaching reduces the efficiency of the method as the emitter density is no longer within the optimal performance window of the NN. Here, we report the integration of DNA-PAINT into image prediction with DeepSTORM, which offers several advantages. First, the concentration of imager strands can be tailored towards obtaining a constant emitter density optimized to the performance window of the NN. Second, generating very low-density emitter data provides true experimental emitters for NN training, which captures the optical properties of the microscope. Third, low-density and high-density emitter data can be generated on the same sample to obtain a ground truth image for each prediction, hence bypassing simulated datasets for NN assessments. Fourth, Exchange-PAINT permits multi-color imaging by exchanging fluorophore-labeled oligonucleotide strands from the imaging buffer, which facilitates multi-target prediction with only a single NN model (Jungmann *et al*., 2014; Narayanasamy *et al*., 2021). Finally, the bleaching-independent nature of DNA-PAINT permits the acquisition of large sample areas in a short time.

Here, we utilize DeepSTORM for the prediction of super-resolution SMLM images from high-density DNA-PAINT data. NN training is performed with experimental low emitter density DNA-PAINT data. Using the trained model, we predict cellular structures in semi-thin neuronal tissue samples with complex structural morphology. Sequential imaging of multiple targets using different oligonucleotides labeled with the same fluorophore enables aberration-free multi-target imaging (Exchange-PAINT) (Jungmann *et al*., 2014) coupled with the use of a single model for multi-color prediction which facilitates the acquisition of information-rich structural data. The image prediction quality was assessed using image-based similarity metrics. In summary, this approach enables data acquisition for an SMLM image within 1 minute for practical multi-color and large-ROI imaging in a fraction of the time compared to conventional multi-color SMLM methods.

## Results

### DeepSTORM model training and prediction workflow

DNA-PAINT is a variant of SMLM that provides a constant signal over time and enables aberration-free multi-color imaging (Schnitzbauer *et al*., 2017). The spatial density of fluorophores in a DNA-PAINT experiment can be easily adjusted by tuning the imager strand concentration in the buffer such that the recording of datasets of the same structure with different fluorophore densities is feasible. These experimental features are ideal for the implementation into neural networks designed to reconstruct SMLM images from high-density single-molecule data (Nehme *et al*., 2018, 2020). To this end, we established a workflow that harnessed the characteristics of DNA-PAINT SMLM to enhance the useability of DeepSTORM. On a whole, very low-density, low-density and high-density emitter data were recorded with DNA-PAINT for model training, ground truth (GT) images, and image prediction, respectively (**Figure 1**). In the first step, we recorded experimental training data at very low emitter density (0.028 emitters/μm^2^) and localized single emitters using the single-molecule localization software Picasso (Schnitzbauer *et al*., 2017) to train a multi-emitter prediction model. This circumvents the need for simulated single-molecule data, and is in line with the report that the prediction performance of DeepSTORM is improved with the use of experimental data for training compared to simulated data (Nehme *et al*., 2018). To generate high-density emitter data for network training, small patches of 16 × 16 pixels with on average one-emitter per frame were generated. These patches were then binned together randomly to output a high-density patch of 2 emitters/μm^2^. These patches, together with the corresponding coordinates of the emitters, were used to train a DeepSTORM model (**Figure 1A**). The trained model was then applied to predict SMLM images from high-density DNA-PAINT data recorded with high concentrations of imager strands (**Figure 1BC**). Concurrently, a single-molecule DNA-PAINT image with low emitter density was generated for the same ROI which served as the GT image (**Figure 1D**). DNA-PAINT data was recorded in semi-thin structurally conserved tissue labeled for α-tubulin and the mitochondrial protein TOM20 (Narayanasamy *et al*., 2021). The predicted images were compared to their respective GT images and the prediction quality was assessed using several quantitative metrics.

**Figure 1:**
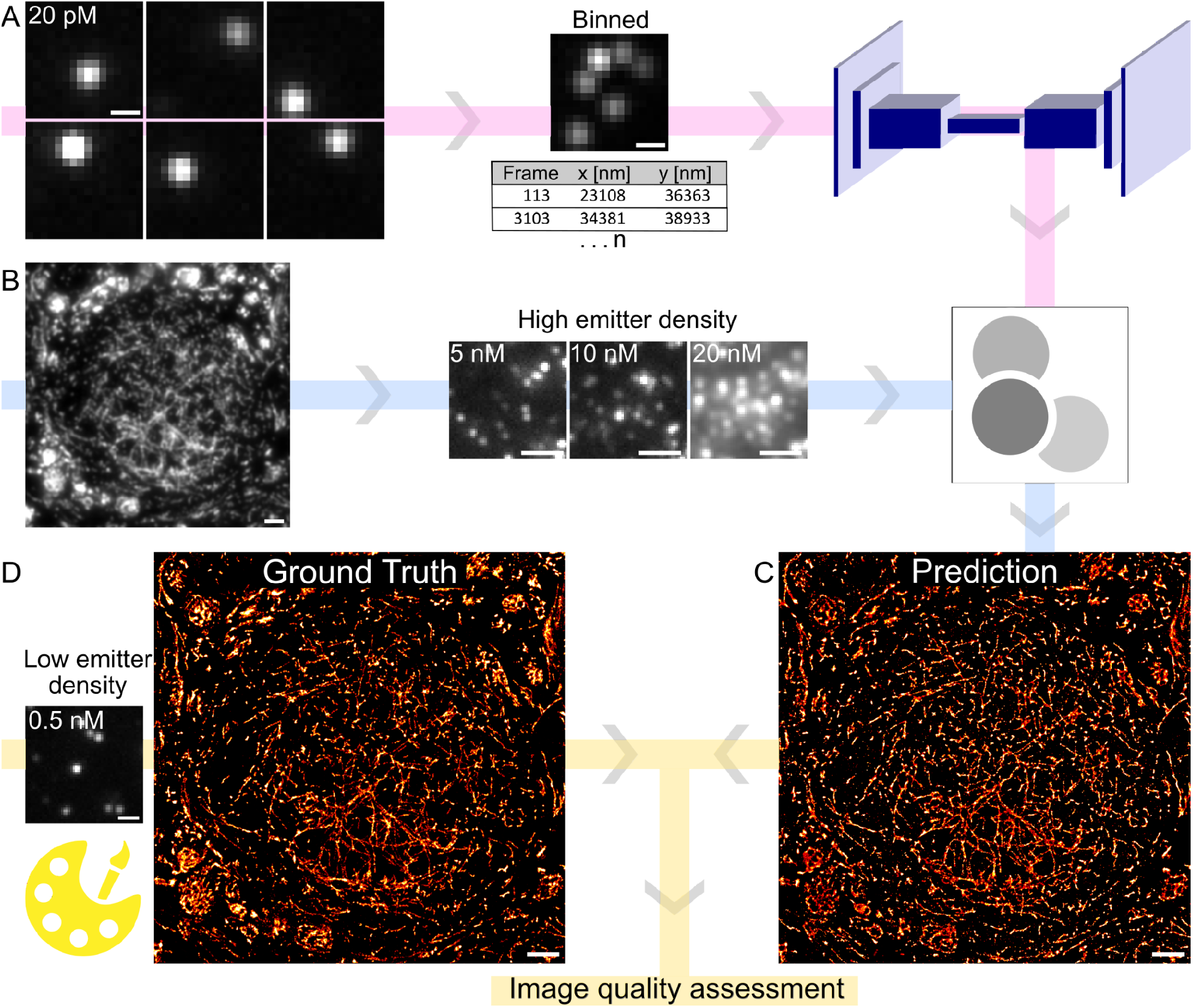
NN-assisted DNA-PAINT imaging. (A) Very low emitter density DNA-PAINT images (20 pM imager strands) were binned into high emitter density images, and together with the single-molecule localizations served as input for the DeepSTORM U-Net architecture to train a DeepSTORM model. (B) DNA-PAINT imaging of tissue samples was performed with different concentrations of fluorophore-labeled imager strands (5, 10, and 20 nM) yielding varying emitter densities. (C) High emitter density DNA-PAINT datasets were input into the trained DeepSTORM and super-resolution images were predicted. (D) For the same sample, a ground truth (GT) image was generated with low emitter density DNA-PAINT (0.5 nM imager strands) and used to assess the quality of the predicted super-resolution image. Scale bars: 0.5 μm (A), 2 μm (B, C & D)

### NN-assisted SMLM imaging in neuronal tissue

We applied the trained model to predict multi-color SMLM super-resolution images. Structurally preserved semi-thin (∼350 nm) cryosectioned rat neuronal tissue sections in the medial nucleus of the trapezoid body (MNTB) region (Klevanski *et al*., 2020) were stained for α-tubulin and TOM20 using DNA-labeled antibodies (see Methods; **Table 1**) and imaged sequentially following the Exchange-PAINT protocol (Jungmann *et al*., 2014; Narayanasamy *et al*., 2021). A super-resolution image reconstructed from low emitter density DNA-PAINT data (0.5 nM imager strands P1, P5; 10000 frames) served as ground truth (**Figure 2A**). For the same sample, high emitter density DNA-PAINT data (5 nM imager strands P1, 10 nM P5; 400 frames) was recorded. Random patches of low-density emitter DNA-PAINT data from an independent sample were binned to create a high-density training dataset based on experimentally recorded emitters on which DeepSTORM was trained. The trained DeepSTORM model was applied to the high emitter density DNA-PAINT data for prediction of the cellular structure (**Figure 2B**). With an integration time of 150 ms, the acquisition of the low emitter density dataset took 25 minutes, whereas the high emitter density dataset took only 1 minute. Visual inspection shows good agreement between GT and predicted super-resolution images, with structures reconstructed faithfully (**Figure 2CD**). The structural features of the five cells (dotted lines in **Figure 2AB**) were predicted and nuclear regions within the cells were clearly defined, as observed in the GT image. Transverse sections of axons (arrow in **Figure 2AB**) and dense circular tubulin bundles in the centre of the image were reproduced in the predicted image. The distribution of mitochondria in the predicted image was correctly reproduced, where mitochondria was found at a higher density within the cytoplasm of cells (**Figure 2AB**).

**Table 1:** Sequences of docking and imager strands.

| Name | Sequence | Modification |
| --- | --- | --- |
| P1 docking strand | TTATACATCTA | 5' - Thiol |
| P5 docking strand | TTTCAATGTAT | 5' - Thiol |
| P1 imager strand | TAGATGTAT | 3' - Cy3B |
| P5 imager strand | CATACATTGA | 3' - Cy3B |

**Figure 2:**
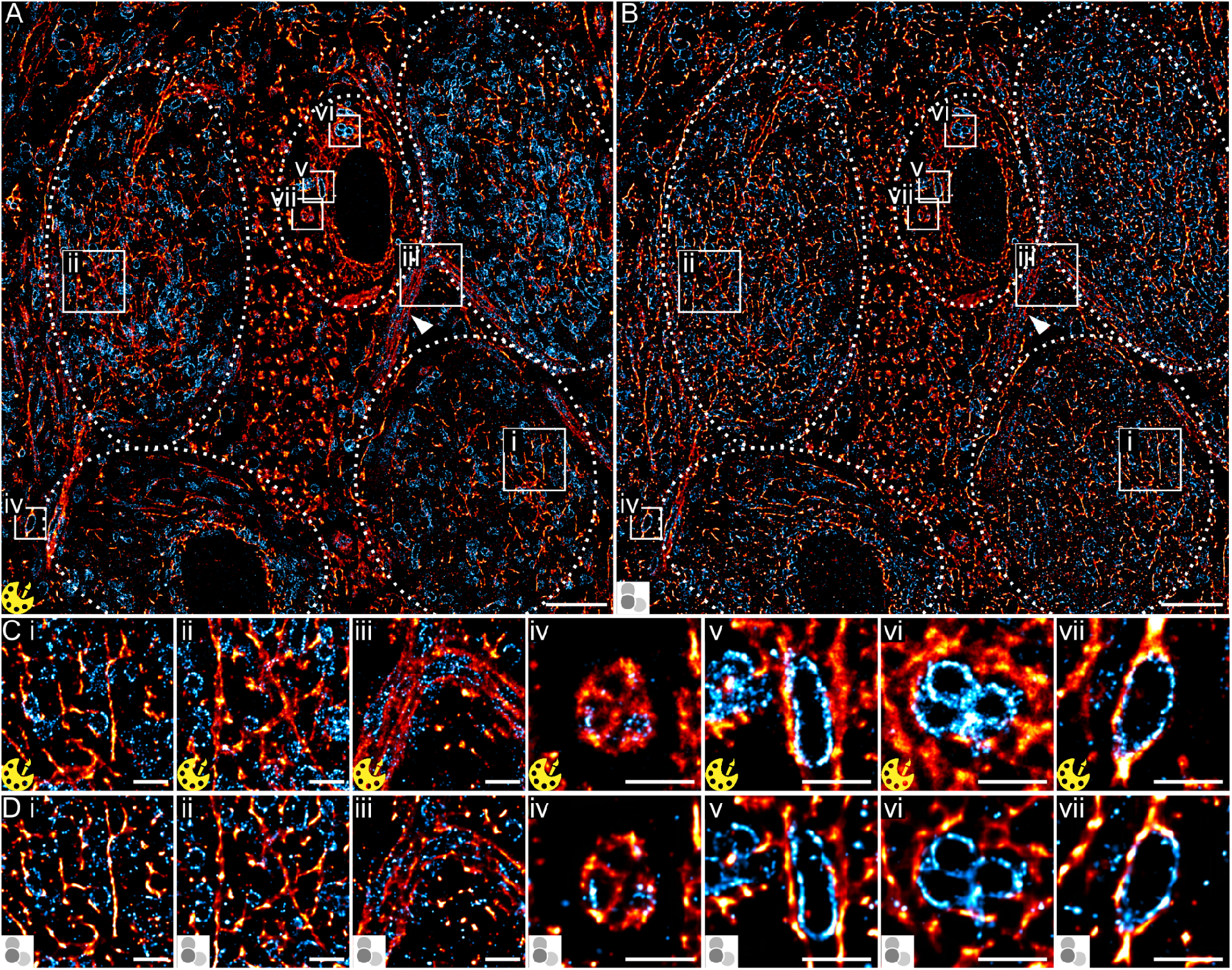
Comparison between Exchange-PAINT super-resolution images and predicted SMLM images in MNTB sections. (A, B) Tissue sample labeled for α-tubulin and TOM20 containing 5 cells (dotted lines) rendered as a (A) GT image (0.5 nM imager strands P1, P5; 10000 frames, 25 minutes acquisition time) and (B) predicted image (5 nM P1, 10 nM P5; 400 frames, 1 minute acquisition time). (C, D) Magnified regions of (C i - vii) GT compared to (D i-vii) predicted images. Scale bars: 5 μm (A & B), 1 μm (C & D i - vii).

To scrutinize the quality of predicted images at a smaller length scale, magnified regions of the experimentally super-resolved structures (GT) (**Figure 2C**) were compared to the predicted structures (**Figure 2D**). Tubulin within the MNTB tissue is found as various morphological structures (Park and Roll-Mecak, 2018; Kelliher, Saunders and Wildonger, 2019), from simple 1-dimensional (1D) to complex 2-dimensional (2D) structures with dense or layered regions. The magnified images show 1D filamentous structures of α-tubulin in the cytoplasm of the principal cell, with thin, elongated, or random patterns that are visually well predicted by the NN (**Figure 2CDi-ii**). Other regions in the tissue show dense and complex 2D arrangements of tubulin (**Figure 2CDiii-vi**) which overall are well predicted in their shape but with reduced performance in their predicted structural density. The structural patterns of TOM20 are mostly uniform and appear as thin, single layer outlines of mitochondria with oblong shapes that can be categorized as 1D structures (**Figure 2Cv-vii**). These structures are predicted very well throughout by the DeepSTORM model, determined by visual inspection and comparison with the GT images of the corresponding mitochondrial regions (**Figure 2CDv-vii**). In summary, we find that our trained DeepSTORM model has excellent prediction quality for the structural features of the two targets labeled in the tissue sections, with a slightly better performance for 1D structures over 2D structures.

### Assessment of image prediction quality

To quantify the quality of SMLM image prediction with the trained DeepSTORM model, we applied image quality metrics and compared the experimental super-resolution data (GT) to the predicted data. First, we applied the HAWKMAN analysis to compare the structural similarity between GT and predicted images (**Figures 3, S1**). HAWKMAN is sensitive to nanoscale differences between images and to artificial sharpening while also providing confidence maps for super-resolved structures (Marsh *et al*., 2021). First, we assessed the quality of structure prediction from high emitter density data for samples stained with TOM20 that were recorded with different imager strand concentrations (5, 10, and 20 nM) (**Figure 3A**). HAWKMAN generated a structure map of skeletonized structures that showed the highest overlap between predicted and GT images for an imager strand concentration of 10 nM (**Figure 3B**, yellow arrows). Similarly, the sharpening map reflects highest structural overlap for an imager strand concentration of 10 nM, whereas at a concentration of 5 nM, the structural envelope of the mitochondria was not completely reconstructed, and at a concentration of 20 nM hallucination artefacts at the edge of structures appeared (**Figure 3C**; arrows). The confidence maps support these findings and show the highest overlap for an imager strand concentration of 10 nM (**Figure 3D**). HAWKMAN analysis of α-tubulin structures in tissue show that structure dimensionality impacts the prediction quality, in that, while 1D structures were predicted well for all three imager strand concentrations, 2D structures were incompletely predicted (**Figure S1**). For further image comparison metrics, we applied (1) SQUIRREL to calculate the resolution-scaled Pearson correlation coefficient (RSP), the resolution-scaled root mean squared error (RSE) and an error map (Culley *et al*., 2018); (2) the multi-scale structural similarity index (MS-SSIM) (Wang, Simoncelli and Bovik, 2003; Prieto, Chevalier and Guibelalde, 2014); and (3) determined the spatial resolution by decorrelation analysis (Descloux, Grußmayer and Radenovic, 2019) (**Figures S2, S3, Supplementary Note 1**).

**Figure 3:**
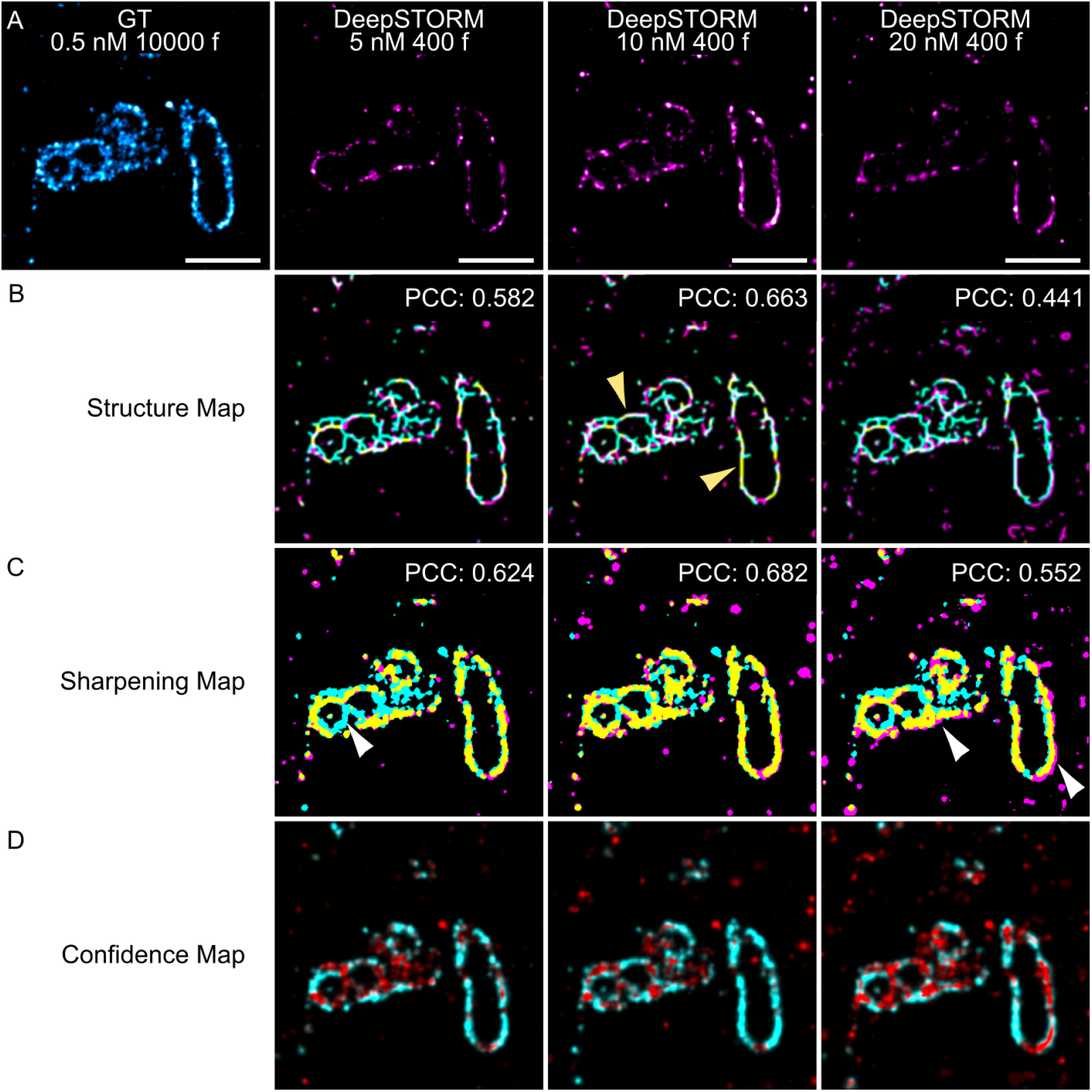
Quantitative analysis of image similarity between ground truth and predicted super-resolution images using HAWKMAN. (A) GT and DeepSTORM predicted images of a TOM20-labeled structure recorded for imager strand concentrations of 5, 10, and 20 nM. (B) Structure map and Pearson correlation coefficient (PCC) indicating regions of good overlap (yellow), denser GT structures (cyan) or denser DeepSTORM predicted structures (magenta). (C) Sharpening map indicating regions of artificial sharpening with the same color scheme as the structure map. (D) Confidence map highlighting structures of high confidence (cyan) and low confidence (red). Scale bars: 1 μm.

### NN-assisted large-ROI super-resolution imaging

The bleaching-independent nature of DNA-PAINT due to the constant replenishment of fluorophore labels enables the recording of large, multi-field-of-view images (Böger *et al*., 2019). We demonstrate this feature in combination with NN-mediated accelerated SMLM imaging of a large-ROI of MNTB tissue, in which calyx of Held synapses are densely organized (**Figure 4A**) (Thomas *et al*., 2019). In a tissue sample labeled for α-tubulin, 16 full-view patches were recorded in 1 minute per image, as opposed to hours when using non-NN DNA-PAINT imaging. This produced a large-view representation of the underlying ultrastructure containing a rich amount of information from the microscale down to the nanoscale (**Figure 4B**). Unlike a confocal image where information breaks down at the nanoscale, or a super-resolution image where only a fraction of cells are found in one image, our stitched multi-patch image possesses a top-down approach where a macroscale overview of a tissue section can be magnified many folds to observe nanoscale details (**Figure S4**). This demonstration shows the potential of NN-assisted, multi-emitter image reconstruction with DeepSTORM for imaging large samples. A straightforward extension to this method is the integration of multiple target labels (**Figure 2**) with multi-patch imaging.

**Figure 4:**
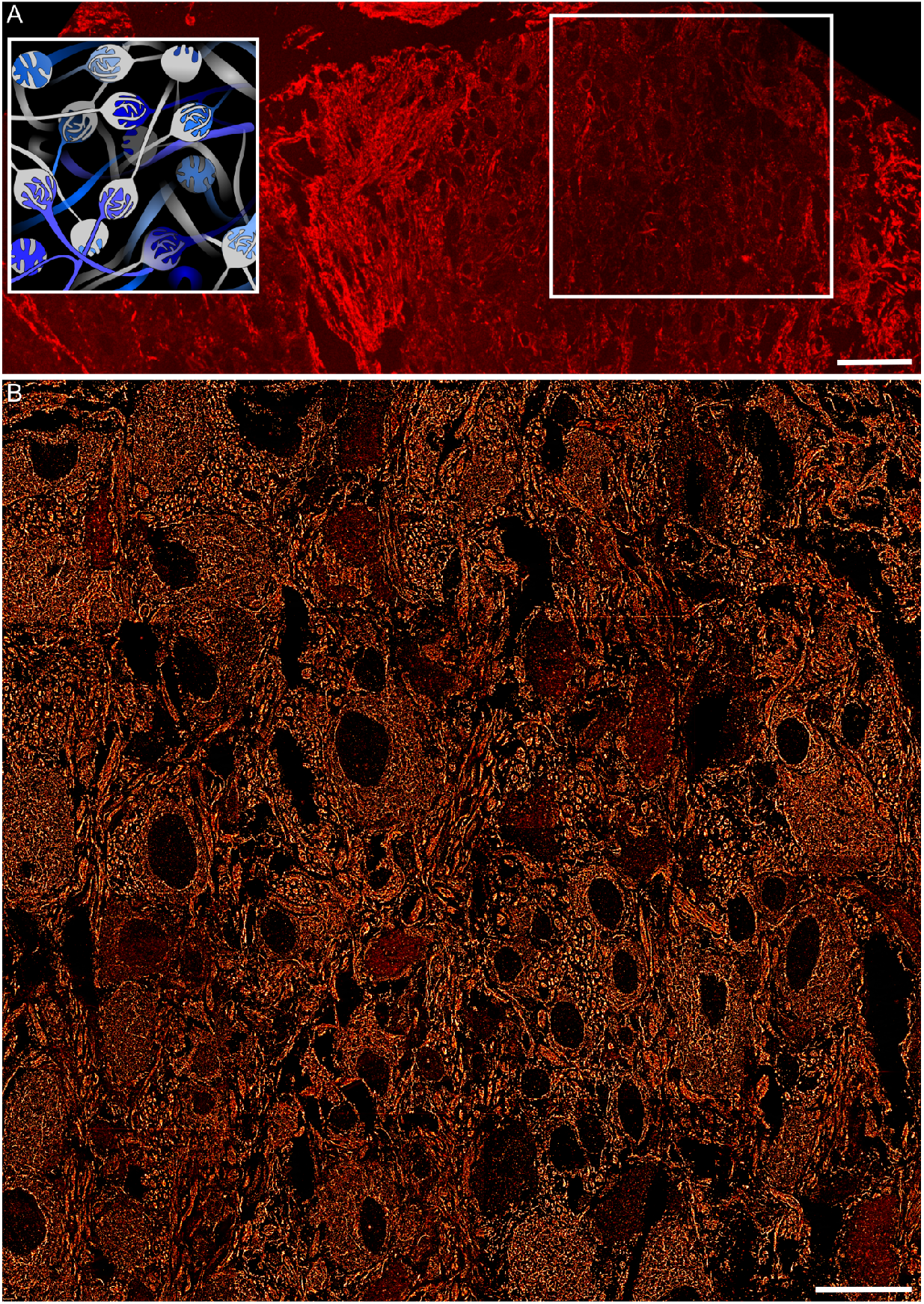
Accelerated large-sample imaging. (A) Confocal microscopy image of an MNTB tissue section and a graphical representation of calyces organized within the MNTB region (inset). (B) Large-ROI super-resolution image recorded for the tissue area defined by the white frame in A. The α-tubulin super-resolution image was obtained by imaging 55 μm x 55 μm patches recorded with 10 nM imager strand P1 in a 4 × 4 grid-like fashion with 400 frames per patch, obtaining high-density DNA-PAINT frames in 1 minute and a total imaging time of 16 minutes. Scale bar 50 μm (A), 20 μm (B).

## Discussion

With the recent developments in artificial intelligence for microscopy, a myriad of tools became available for SMLM, and with this the challenge of optimizing the interface between image data and computational treatment (Laine *et al*., 2021). Here, we present an experimental workflow that facilitates the use of neural networks for high emitter density image prediction by introducing the unique imaging features of DNA-PAINT. The complementarity with DNA-PAINT imaging makes the application of these networks more robust, extends their capabilities and removes barriers for their everyday implementation. Key features to this method are (1) a constant and adjustable emitter density over time, maneuvering the experimental data into the optimal performance window of a NN; (2) NN training with experimental imaging data without the need for simulated single-molecule data; (3) sequential imaging rounds of the same sample, which facilitate the recording of low and high density data from the same structure for robust quantitative image quality assessment; (4) multi-target prediction with only a single NN-trained model for various structures; (5) large-ROI imaging by the sequential imaging of multiple regions within a large sample.

We implemented these experimental features and demonstrated NN-assisted prediction of super-resolved cellular structures in structure-conserved semi-thin brain tissue, using the DeepSTORM network (Nehme *et al*., 2018). Key advantages to using DeepSTORM are (1) its significant acceleration in image acquisition time, (2) reduced drift due to short image acquisition time which in turn improves localization precision (Costello and Cox, 2021), and (3) the reduced need for data storage capacity. In this study, a 1-minute imaging time at 5 - 10 nM imager strand concentration was sufficient to produce structures comparable to GT images. Previous studies have compared DeepSTORM prediction to leading multi-emitter algorithms and found that DeepSTORM computed much faster and with better accuracy to ThunderSTORM (von Chamier *et al*., 2021), FALCON and CEL0 (Nehme *et al*., 2018). Furthermore, DeepSTORM is structure independent in that one model can be used for predicting various targets/structures without generating hallucination artefacts stemming from memorizing structural features. For the implementation of Exchange-PAINT, only a single model is sufficient for predicting multiple experimental structures, thereby further generalizing and improving the accessibility of the method. Of note, the workflow we demonstrated is compatible with other high-emitter NNs. Previous studies evaluated the performance of DeepSTORM in simulated and experimental data using different analysis metrics (Nehme *et al*., 2018; von Chamier *et al*., 2021). Other studies used DeepSTORM as a benchmark to assess the performance of novel dense-emitter NNs (Dardikman-Yoffe and Eldar, 2020; Yao *et al*., 2020). To establish uniformity in analyses for similar studies, we propose several tools that quantify image similarity to be used to assess the performance of SMLM-based DL tools. We found that SQUIRREL (Culley *et al*., 2018) and HAWKMAN (Marsh *et al*., 2021) are complementary analysis methods, where the former expounds intensity discrepancies whereas the latter focuses on nanoscale structural (dis-)similarities. We also note that other tools for quantitative image comparison are available (Sage *et al*., 2019; Chen and Chen, 2021; Speiser *et al*., 2021). We found that a combination of visual checks and quality metrics were most suitable for assessing prediction quality (**Figures 3, S2 and S3**).

Further to image similarity, spatial resolution is a relevant parameter in predicted and GT images. We applied decorrelation analysis (Descloux, Grußmayer and Radenovic, 2019) and found that the spatial resolution in predicted images was, throughout all imaging conditions (5, 10, and 20 nM imager strands), slightly higher (∼ 45 nm) than in GT images (∼ 35 nm) (**Figure S2E**). The difference in spatial resolution could be attributed to a number of reasons such as the method of rendering by different software, the effect of structure dimensionality, or the local density of emitters which may impair the quality of a predicted image (**Figure 2CD**).

Comparisons between experimental and simulated PSFs showed that DeepSTORM trained on experimental PSFs had better prediction precision (Nehme *et al*., 2018). To utilize this to our advantage, the training dataset used for our model was derived from experimental PSFs on the same optical setup (**Figure 1**). The advantage of DNA-PAINT is evident here as the imager strand concentration can be reduced until a sparse emitter dataset is obtained, suitable for isolating single-PSF patches. This further reduces the need for parameter analysis for artificial PSF generation and reduces the PSF error margin between training patches and prediction datasets.

The performance of a model trained with DeepSTORM has an optimal operation range with respect to emitter densities, and image prediction might break down above a certain density threshold. Nehme et al. report good network performance up to 6 emitters/μm^2^ (Nehme *et al*., 2018). In this work, a range of imager stand concentrations were used to determine the best prediction output. Increasing the imager strand concentration results in an increase in emitter density, which reduces the number of frames required to obtain a fully formed image, hence improving temporal resolution. However, beyond this point, one introduces (1) too high emitter densities which are then predicted with lower accuracy and yield worse spatial resolution, and (2) higher fluorescence background in the buffer, which reduces frame signal-to-noise ratios (SNRs), to which DeepSTORM is susceptible (Nehme *et al*., 2018). These tradeoffs are to be considered when choosing the right imager strand concentration. The prediction quality is also dependent on the dimensionality of structures where complex 2D shapes were reconstructed with lower precision compared to simple 1D structures (**Figure 2**). Consequently, we found that an optimal imager strand concentration is structure dependent, with dense structures like tubulin requiring lower concentrations compared to mitochondria. Using Exchange-PAINT, the optimal density of emitters can be tailored towards the structures being imaged, thereby maintaining good image quality and short imaging time. Nevertheless, DeepSTORM prediction was found to be very robust as the model was able to handle a range of emitter densities, from 5 to 10 nM imager strand concentrations (**Figures 3, S2 and S3**). At 20 nM, DeepSTORM performance deteriorated, likely due to lower SNR and locally excessively overlapping emitters. High emitter density is not only an issue in high-density DL tools but also in conventional SMLM methods where the reconstructed image appears sharp or smooth and contains artefacts (Costello and Cox, 2021). The blob-like appearance of predicted images is also a feature of DeepSTORM, which becomes more evident at very high imager strand concentrations.

In conclusion, the combination of DNA-PAINT SMLM with a multi-emitter NN has proven to be a robust method for super-resolution structure prediction in neuronal tissue. The model was able to generalize well for a range of emitter densities. Furthermore, the concurrent use of DNA-PAINT and DeepSTORM allows for more control over emitter densities and further enhances DeepSTORM efficiency as the whole dataset is at its optimal working range. With the constant emitter density and photostability of DNA-PAINT, a large-ROI can be imaged in a matter of minutes. Before the incorporation of DL tools into super-resolution microscopy, there had been a tradeoff between image size and image resolution. Based on the proof-of-concept shown here, it is possible to overcome this tradeoff using DL tools to be able to get a bird’s eye view of the sample while also magnifying down to the nanoscopic details of individual proteins. This, coupled with Exchange-PAINT to visualize multiple protein targets that can be predicted with a single model, will develop into a powerful tool for biomedical imaging.

## Methods

### Tissue preparation

All experiments that involved the use of animals were performed in compliance with the relevant laws and institutional guidelines of Baden–Württemberg, Germany (protocol G-75/15). Animals were kept under environmentally controlled conditions in the absence of pathogens and *ad libitum* access to food and water. Preparation of brain sections containing the MNTB was performed according to an established protocol (Klevanski *et al*., 2020) with slight modifications. Briefly, Sprague-Dawley rats (Charles River) at postnatal day 13 were anaesthetized and perfused transcardially with PBS followed by 4% PFA (Sigma-Aldrich). Brains were dissected and further fixed in 4% PFA overnight at 4°C. The next day, 200 μm thick vibratome (SLICER HR2, Sigmann-Elektronik, Germany) sections of the brainstem containing MNTB were prepared. MNTB were excised and infiltrated in 2.1 M sucrose (Sigma-Aldrich) in 0.1 M cacodylate buffer overnight at 4°C. Tissue was mounted on a holder, plunge-frozen in liquid nitrogen in 2.1 M sucrose and semi-thin sections (350 nm) were cut using the cryo ultramicrotome (UC6, Leica). Sections were picked up with a custom made metal loop in a droplet of 1% methylcellulose and 1.15 M sucrose and transferred to 35 mm glass bottom dishes (MatTek, USA) pre-coated with 30 μg/ml of fibronectin from human plasma (Sigma-Aldrich) and nanodiamonds (100 nm; Adamas Nanotechnologies, USA) as fiducials. Dishes containing sections were stored at 4°C prior to their use.

### Antibody-DNA conjugation

Secondary antibodies of donkey anti-mouse (715-005-151) and donkey anti-rabbit (711-005-152) were purchased from Jackson ImmunoResearch. DNA strands were purchased from Metabion with a thiol modification on the 5′ end for each docking strand and a Cy3B dye on the 3′ end for the imager strands (**Table 1**).The antibody to DNA docking strand conjugation was prepared using a maleimide linker as previously reported in detail (Schnitzbauer *et al*., 2017). The thiolated DNA strands were reduced using 250 mM DTT (A39255, ThermoFisher Scientific). The reduced DNA was purified using a Nap-5 column (17085301, GE Healthcare) to remove DTT and concentrated with a 3 kDa Amicon spin column (UFC500396, Merck Milipore). Antibodies (>1.5 mg/mL) were reacted with the maleimide-PEG2-succinimidyl ester crosslinker in a 1:10 molar ratio, purified with 7K cutoff Zeba desalting spin columns (89882, ThermoFisher Scientific) and concentrated to >1.5 mg/mL.The DNA and antibody solutions were cross-reacted at a 10:1 molar ratio overnight and excess DNA was filtered through a 100 kDa Amicon spin column (UFC510096, Merck Milipore). The antibody-DNA solution was stored at 4°C.

### Tissue labeling

Tissue samples were labeled with primary antibodies against α-tubulin-mouse (T6199, Sigma-Aldrich) and TOM20-rabbit (sc-11415, Santa Cruz Biotechnology). Tissue samples in dishes were washed with PBS three times for 10 min each to remove the sucrose-methylcellulose layer and blocked with 5% fetal calf serum (FCS) for 30 min. The primary antibodies were diluted in 0.5% FCS and applied to the tissue section for 1 h at room temperature (rt) and washed off three times with PBS. The conjugated secondary antibody-DNA docking strand in 0.5% FCS was applied onto tissue for 1 h at rt and washed 3 times with PBS. The tissue was then stained with Alexa Fluor 488-conjugated WGA (WGA-A488) (W11261, Thermo Fisher Scientific) in PBS for 10 min and washed off three times with PBS.

### SMLM setup

DNA-PAINT microscopy was performed on a home-built SMLM setup with an Olympus IX81 inverted microscope frame equipped with an Olympus 150x TIRF oil immersion objective (UIS2, 1.49NA). The samples were illuminated in TIRF mode using a 561 nm laser line (Coherent Sapphire LP) at an illumination density of 0.88 kW/cm^2^ through a 4L TIRF filter (TRF89902-EM, Chroma Technology) and ET605/70 M nm bandpass filter (Chroma Technology). Signals were detected with an Andor iXon EM+ DU-897 EMCCD camera (Andor, Ireland). SMLM frames were acquired using multi-dimensional acquisition (MDA) mode in Micro-Manager 2.0 (Edelstein *et al*., 2014).

### DNA-PAINT imaging

DNA-PAINT imaging was performed in Buffer C (2.5 M NaCl; S7653, Sigma-Aldrich in 5x PBS; 14200-059, Gibco Fisher Scientific) supplemented with 1 mM ethylenediaminetetraacetic acid (EDTA; E6758, Sigma-Aldrich), 2.5 mM 3,4-dihydroxybenzoic acid (PCA; 03930590, Sigma-Aldrich), 10 nM protocatechuate 3,4-dioxygenase pseudomonas (PCD; P8279, Sigma-Aldrich), and 1 mM (±)-6-hydroxy-2,5,7,8-tetramethylchromane-2-carboxylic acid (Trolox; 238813-5G, Sigma-Aldrich). To obtain images for training the DeepSTORM model, 20 pM P5 imager strands were imaged in TOM20 labeled tissue samples. For conventional DNA-PAINT imaging with Picasso software (v0.2.8) analysis to obtain a GT super-resolution image, P strands (P1 and P5) were imaged at an imager strand concentration of 0.5 nM for 10000 frames and acquisition rate of 150 ms for both α-tubulin and TOM20. High-density emitter DNA-PAINT datasets for DeepSTORM image prediction were obtained by imaging α-tubulin and TOM20 at imager strand concentrations of 5 nM, 10 nM, and 20 nM for 400 frames. Exchange-PAINT was performed manually by adding the imaging buffer to the sample chamber and acquiring camera images. The buffer was then removed and the sample washed five times with 1× PBS to remove all imager strands. The subsequent imaging buffer containing the second imager strand was then added and the procedure repeated to image the second target.

Raw DNA-PAINT frames imaged with 0.5 nM imager strands were processed and rendered using Picasso software (Schnitzbauer *et al*., 2017). Events in each frame were localized by fitting using the Maximum Likelihood Estimation for Integrated Gaussian parameters (Smith *et al*., 2010). The localized events were then filtered by their width and height of the Point Spread Function (sx, sy). The resulting localizations were drift-corrected using redundant cross-correlation (RCC), rendered using the ‘One Pixel Blur’ function and further processed using the ‘linked localizations’ function to merge localizations that appeared in multiple consecutive frames. Rendered images were oversampled to match the pixel size of DeepSTORM images. Images were merged in Fiji (Schindelin *et al*., 2012) using the ‘merge channels’ tool and aligned by linear transformation using nanodiamonds as registration reference. The individual channels were assigned pseudocolors.

Super-resolution large sample imaging on α-tubulin was performed using DNA-PAINT imaging with 10 nM P1 imager strands. Four hundred DNA-PAINT frames per imaging area were acquired in a grid-like fashion of 4×4 with an overlap of ∼10% between images. The images were registered using Inkscape software based on structural similarity. The whole image is available on https://doi.org/10.5281/zenodo.5576100. Confocal microscopy for α-tubulin was performed on a Nikon C2 Plus with a Nikon Plan Fluor 40x oil immersion objective (NA 1.30). The tissue sample was imaged on 300 nM P1 imager strands in Buffer C 1x with a 561 nm excitation laser.

### Image binning

DeepSTORM model training required an artificially binned dataset generated from experimental DNA-PAINT data. This data set contains overlapping point spread functions of single emitters together with their precise localization coordinates. A custom script was written for this task and is available at https://github.com/JohannaRahm/ImageBinner (*ImageBinner* version 210408, Python 3.9.2). Patches from the sparse frames and their localization coordinates from Picasso localization software were randomly selected and merged to create high-density emitter patches with matching localization lists. The binning of *n* number of patches introduces camera noise which was corrected by subtracting the value of the camera noise *n-1* times from the high-emitter density patches. The camera noise was estimated as the average pixel intensity of frames acquired with a closed shutter.

A low emitter density DNA-PAINT dataset of tissue labeled for TOM20 was recorded using an imager strand concentration of 20 pM to obtain sparse and isolated single events at a density of 0.028 emitters/μm^2^. To generate training patches, 5000 DNA-PAINT frames of 512 × 512 pixels were input into the *Image Binner* software. A minimum of 1 emitter per patch (17 × 17 pixels) was produced. These patches were binned randomly to generate 30000 high-density emitter patches at a mean emitter density of 1.9 emitters/μm^2^ with a 17 × 17 pixel patch size and its corresponding localization coordinate list.

### DeepSTORM training and prediction

DeepSTORM model training was performed on Google Colab. The resources allocated for DeepSTORM on Colab was NVIDIA-SMI 460.56 with CUDA version 11.2 and Tensorflow version 2.4.1 or 2.5.0. The model used for prediction was trained with 30000 binned patches and a density of 1.9 emitter/μm^2^. Training took 35 minutes with ColabPro.

Raw images with low emitter density, high emitter density binned patches used for NN training, and model metadata are available at https://doi.org/10.5281/zenodo.5704569. Binned image patches along with the localization list served as input for the ZeroCostDL4Mic Colab notebook (von Chamier *et al*., 2021). To directly use the binned image patches as input, the number of patches per frame was set to 1 and the patch size to 16. The maximum number of patches was set to 30000, minimum number of patches to 1, and default values were used for other parameters. Training parameters were set with a number of epochs of 100, batch size of 256, number of steps of 0, percentage validation of 15, and initial learning rate of 10^−5^.

For high emitter density image prediction, 512 × 512 pixels of 400 frames were input into DeepSTORM. A batch size of 1 was used with default values for other parameters. Predictions were performed on DNA-PAINT frames with imager strand concentrations of 5 nM, 10 nM, and 20 nM. Prediction took 7 to 25 minutes depending on the resources allocated by Colab (Colab/ColabPro).

### Image analysis

Picasso-rendered ground truth (GT) and DeepSTORM predicted super-resolution images were visualized and analyzed in Fiji (Schindelin *et al*., 2012). The spatial resolution was calculated for GT and DeepSTORM predicted images using an ImageJ plugin for decorrelation analysis (Descloux, Grußmayer and Radenovic, 2019). For each target (α-tubulin and TOM20), GT image and the three predicted images were merged and registered using the Register Channels tool in the NanoJ Core plugin (Descloux, Grußmayer and Radenovic, 2019; Laine *et al*., 2019). Multi-scale structural similarity index was measured using the MS-SSIM plugin in Fiji (Wang, Simoncelli and Bovik, 2003; Prieto, Chevalier and Guibelalde, 2014). GT and predicted images were intensity-normalized and registered. Each predicted image was compared to GT to obtain the multi-scale structural similarity index between two images. Exchange-PAINT images (α-tubulin and TOM20 in a single ROI) for GT were registered using fiducial markers and DeepSTORM-Exchange-PAINT images were registered with GT as a reference.

For SQUIRREL analysis (Culley *et al*., 2018), predicted images (with imager strand concentrations of 5 nM, 10 nM, 20 nM) were used as reference images against GT images (with imager strand concentration of 0.5 nM, rendered with Picasso) as the test images. A magnification factor of 1 was used. The GT images were intensity-normalized and Point Spread Function (PSF) convolved by SQUIRREL, and an error map, Resolution Scaled Error (RSE), and Resolution Scaled Pearson (RSP) was output. SQUIRREL was also used to compare GT images with diffraction-limited images. Raw DNA-PAINT frames were z-projected with ‘average intensity’ for the number of frames used to render the final super-resolution image, i.e. 10000 frames for 0.5 nM GT image, and 400 frames for the 5, 10, and 20 nM DeepSTORM predicted images. The z-projected image was input into SQUIRREL as a reference image and compared to its corresponding super-resolution image, yielding an error map, RSE, and RSP as output. For HAWKMAN analysis (Marsh *et al*., 2021), super-resolution GT and predicted images were registered and converted to 8-bit. The images were input into HAWKMAN with GT as reference images and DeepSTORM prediction as test images. The calculated length scale was 23 nm/pixel. A pixel scale of 3 corresponding to a 69 nm length scale was chosen for the analysis based on the upper bounds of decorrelation resolution of predicted images.

## Funding

MH acknowledges the funding by the Baden-Württemberg Foundation (Mult!Nano, Methods in life sciences program), in whose name this research was conducted. MH and JR acknowledge funding by the Deutsche Forschungsgemeinschaft (DFG, German Research Foundation) – Project number 414985841, GRK 2566.

## Acknowledgements

We are grateful to Ulrike Engel from the Nikon Imaging Centre, Heidelberg University for assistance with confocal microscopy, Carlo Beretta for implementation of DeepSTORM on a local machine, and Maja Klevanski, Alekasandar Stojic and Thomas Kuner for animal tissue samples. We thank Christoph Spahn for SQUIRREL assistance, Marina Dietz and Yunqing Li for providing DNA-PAINT docking strands. We are grateful to Elias Nehme for advice and assistance with DeepSTORM.

## Notes

https://doi.org/10.5281/zenodo.5576100

https://github.com/JohannaRahm/ImageBinner

### Competing Interest Statement

The authors have declared no competing interest.

