## Supplementary Information for "Fast DNA-PAINT imaging using a deep neural network"

Kaarjel K. Narayanasamy[1,2], Johanna V. Rahm[2], Siddharth Tourani[3], Mike Heilemann[1,2]
[1] Department of Functional Neuroanatomy, Institute for Anatomy and Cell Biology, Heidelberg University, Heidelberg, Germany
[2] Institute of Physical and Theoretical Chemistry, Goethe University Frankfurt, Frankfurt, Germany
[3] Visual Learning Lab, Heidelberg University, Heidelberg, Germany

### Supplementary Information

##### Supplementary Note 1

An α-tubulin-labeled region in tissue with 1D (left) and 2D (right) structures in the same image are compared (**Figure S1A**). The structure map shows that 1D α-tubulin filaments are predicted well by DeepSTORM but the 2D α-tubulin bundles are lacking structural density in the predicted images (**Figure S1B**). Similarly, stronger artificial sharpening (dense cyan regions) are observed at the 2D regions compared to the 1D tubulin (**Figure S1C**). The confidence maps corroborate these findings where low reconstruction correlation in the 2D regions and good correlation in the 1D regions are evident (**Figure S1D**).

Image prediction quality was assessed for α-tubulin- and TOM20- labeled images using HAWKMAN, SQUIRREL, MS-SSIM, and decorrelation resolution (**Figure S2**). The overlay between GT and predicted images (5, 10, or 20 nM) indicate either structural agreement (white), denser GT structures (cyan) or denser predicted structures (magenta) (**Figure S2A**). Visual comparisons indicate 5 nM and 10 nM predictions of α-tubulin are highly similar (**Figure S2Ai-ii**) whereas a noticeable difference is observed in 20 nM where the cyan in structurally dense regions are more prominent ((**Figure S2Aiii**; arrows). Here, DeepSTORM loses its prediction quality at 20 nM for dense 2D α-tubulin structures while maintaining the reconstruction of 1D structures. The comparison of a dense 2D structural region of an axon (Narayanasamy *et al.*, 2021) for GT (cyan) and DeepSTORM 5, 10 and 20 nM imager strand concentration (magenta) was assessed (**Figure S2B**). The prediction quality for 2D structures is low, exacerbated by increasing imager strand concentrations (**Figure S2Bi-iv**). The SQUIRREL error maps indicate larger errors with increasing concentration of imager strands and dissimilarities in structures can be seen in blue and green regions, while yellow regions indicate differences in intensity (**Figure S2Bv-vii**).

Visual inspection of TOM20 predicted images compared to GT indicate that an imager strand concentration of 5 nM is not sufficient to completely reproduce mitochondrial structures (high cyan density), whereas at 10 nM there is better similarity between GT and the predicted image (**Figure S2Ci-ii**). At 20 nM hallucination artefacts are being predicted in the DeepSTORM image

which are not found in GT (**Figure S2Ciii**). A magnified region of a single mitochondria shows that the GT image is finer and more punctate compared to the larger and diffuse points of the predicted images (**Figure S2Di-iv**). While the mitochondrial structure and shape were effectively reconstructed in all three predicted imager strand concentrations, the 5 nM imager strand prediction is incomplete (arrows). At 10 nM, the mitochondrial shape is more defined and better reproduced, and at 20 nM hallucination artefacts are formed along the structural boundary (arrows). The error maps show very subtly that 10 nM imager strand concentration has the lowest structural error. Strong yellow regions dotted around the structure reflect differences in intensity rather than structural inconsistencies (**Figure S2Dv-vii**; arrows) (Marsh *et al.*, 2021).
The quality of DeepSTORM predicted structures compared to GT were quantitatively assessed (**Figure S2E**). For SQUIRREL analysis, α-tubulin showed slightly higher RSP values for 5 nM imager strand concentrations while no difference was observed for TOM20 RSP values in all imager strand concentrations. α-tubulin and TOM20 both have the lowest RSE at 5 nM (p α-tubulin = 0.02; ANOVA). This suggests that an increase in imager strand concentration contributes to higher background fluorescence which affects image prediction quality. The MS-SSIM for α-tubulin had the highest structural similarity at 5 nM imager strand whereas in TOM20-labeled structures only 20 nM was unsuitable for prediction. In the HAWKMAN analysis for both structural reconstruction and artificial sharpening, α-tubulin at 5 nM performed well ($p = 0.02$; ANOVA) whereas TOM20 had comparable structural correlation for 5 and 10 nM and better sharpening correlation at 10 nM. Decorrelation resolution for both α-tubulin and TOM20 are lower in all predicted images compared to their respective GT images (~35 nm) by approximately 10 nm.

We applied SQUIRREL analysis on the super-resolved low-density emitter and high-density predicted DeepSTORM images against their respective diffraction-limited DNA-PAINT frames obtained by z-projection. The low-density emitter image showed the best RSP value and the lowest RSE compared to DeepSTORM predicted images, also reflected in the error map showing high image correlation (**Figure S3Aiii**). With the increase in imager strand concentrations for predicted images, the RSP, RSE, and error map become worse. While the 1D structures on the left side of the images are largely unchanged in the error map, the prediction quality of 2D dense structures (right side) become noticeably poor (**Figure S3Aiv-vi**). Similar to α-tubulin, the low-density emitter image for TOM20 showed the best outcome with the highest RSP and lowest RSE value compared to predicted images (**Figure S3Biii-vi**). Again, both the RSP and RSE values suffered with increasing imager strand concentrations, although the difference between 5 and 10 nM was low in mitochondrial structures. The reduced prediction quality with increasing imager strand concentrations may be attributed to excessive overlap of emitters and high background fluorescence. The prediction is also affected by structure dimensionality, whereby 1D structures were predicted better than 2D structures.

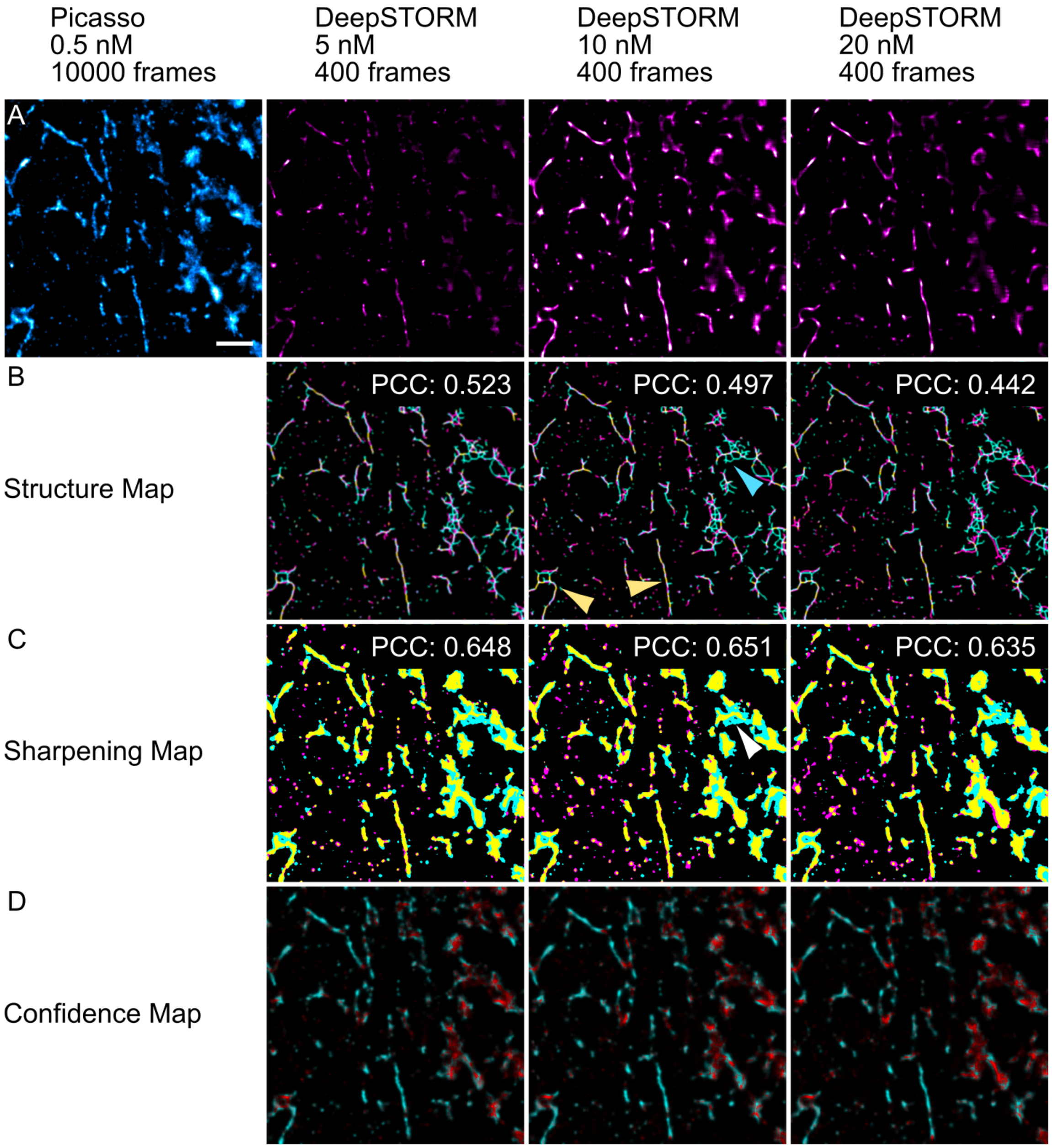


**Figure S1**: **Quantitative analysis of image similarity between ground truth and predicted super-resolution images using HAWKMAN.** HAWKMAN analysis of a region of α-tubulin between GT and predicted images. (A) GT and DeepSTORM predicted images of an α-tubulin-labeled structure recorded for imager strand concentrations of 5, 10, and 20 nM. (B) Structure map and Pearson correlation coefficient (PCC) indicating regions of good overlap (yellow), denser GT structures (cyan) or denser DeepSTORM predicted structures (magenta). (C) Sharpening map indicating regions of artificial sharpening with the same color scheme as the structure map. (D) Confidence map highlighting structures of high confidence (cyan) and low confidence (red). Scale bars: 1 μm.

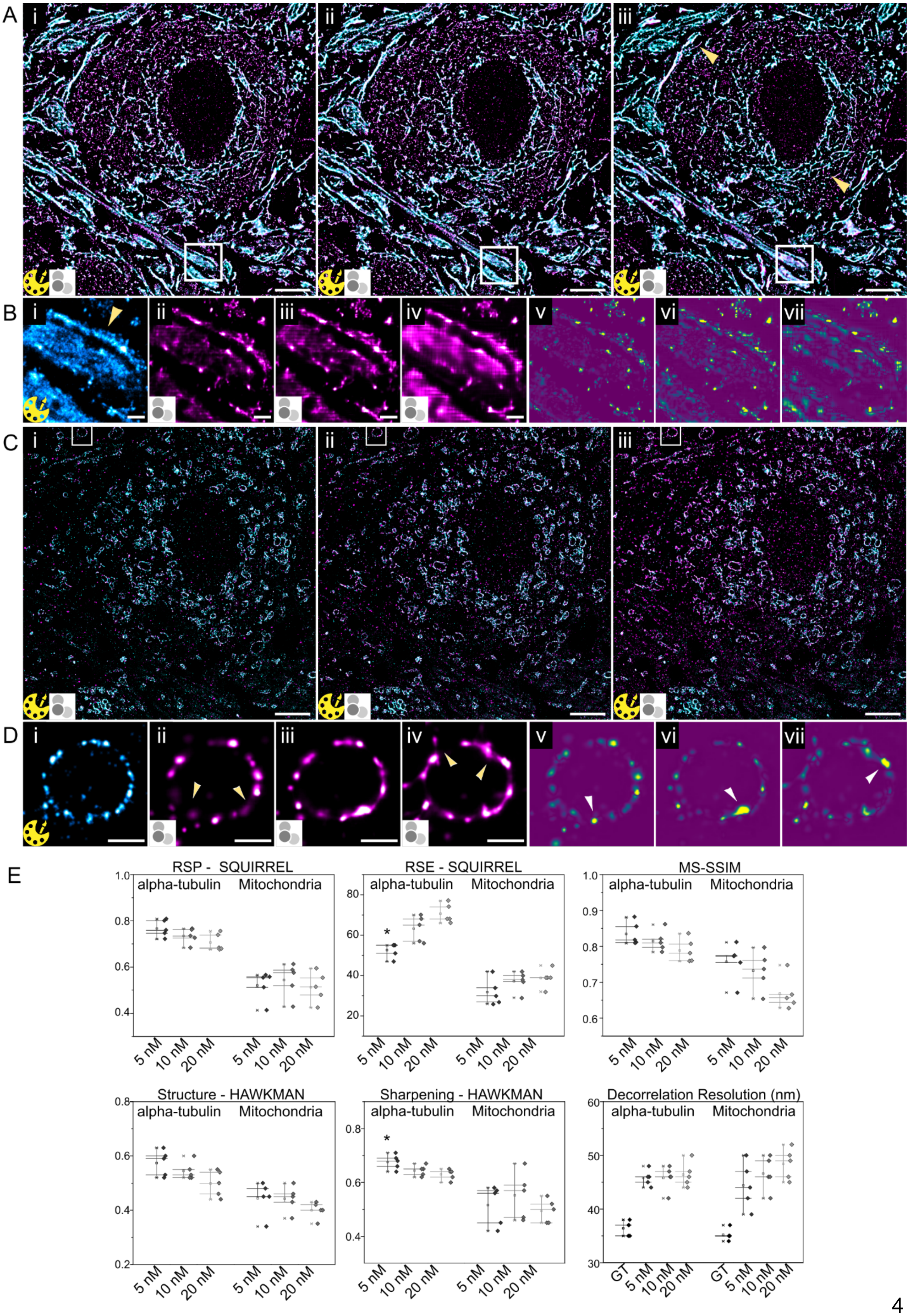
A
i
ii
iii
B
i
ii
iii
iv
v
vi
vii
C
i
ii
iii
D
i
ii
iii
iv
v
vi
vii
E
RSP - SQUIRREL
alpha-tubulin
Mitochondria
5 nM
10 nM
20 nM
RSE - SQUIRREL
MS-SSIM
Structure - HAWKMAN
Sharpening - HAWKMAN
Decorrelation Resolution (nm)
GT

**Figure S2: Quantitative assessment tools for the analysis of predicted super-resolution images.** (A) Overlay of GT α-tubulin-labeled images (0.5 nM, 10000 frames; cyan) with (i) 5 nM, (ii) 10 nM, and (iii) 20 nM DeepSTORM predicted images (400 frames; magenta). (B) A magnified region from A with (i) GT; cyan, (ii) 5 nM, (iii) 10 nM, and (iv) 20 nM predicted images (magenta). SQUIRREL error map between GT and (v) 5 nM, (vi) 10 nM, and (vii) 20 nM predicted images. (C) Same as A for TOM20. (D) Same as B for TOM20. (E) Predicted image quality assessment against GT image using RSP (Resolution Scaled Pearson) and RSE (Resolution Scaled Root Mean Squared Error) from SQUIRREL; MS-SSIM (Multi-Scale Structural Similarity Index); Pearson Correlation-based (PCC) structure similarity and image sharpening with HAWKMAN; and decorrelation resolution; n=5. Scale bar 4 µm (A & C), 0.5 µm (B & D).

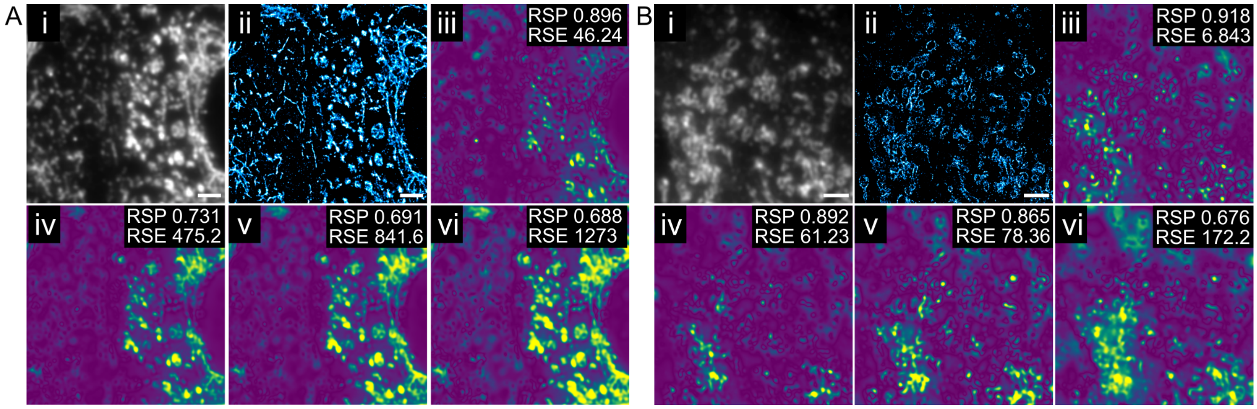


**Figure S3: SQUIRREL analysis between super-resolution and diffraction-limited images** (A) α-tubulin-labeled structure with (i) diffraction-limited image of 0.5 nM imager strands for 10000 frames and (ii) corresponding low-density emitter super-resolution image. SQUIRREL analysis (iii) between ii and i, and between DeepSTORM predicted images and diffraction-limited images (400 frames) for (iv) 5 nM (v) 10 nM and (vi) 20 nM imager strands (B) Same as A for TOM20-labeled structures. Scale bar 2 µm.

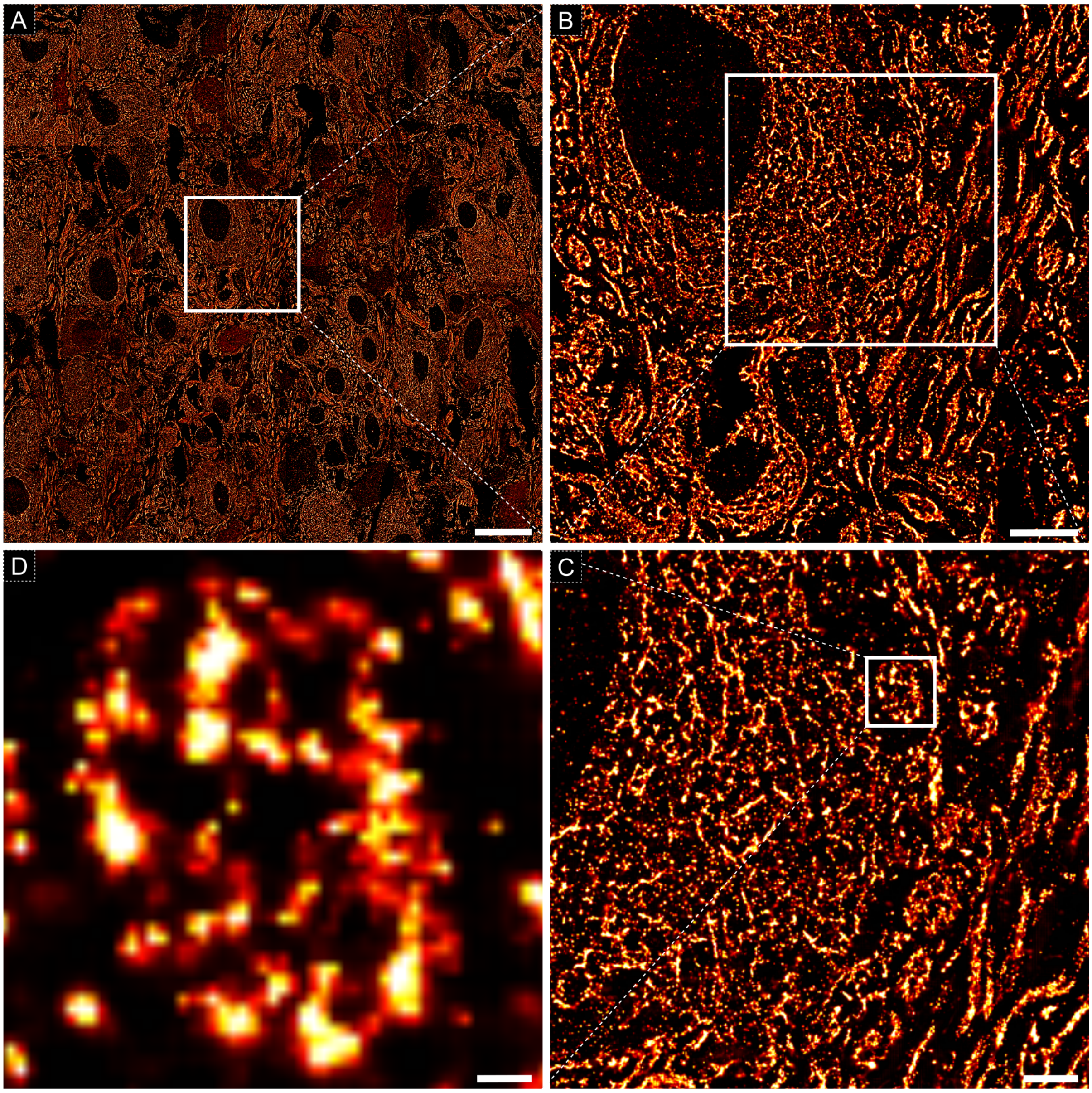


**Figure S4: Magnified super-resolution large-sample image.** (A) A large-ROI super-resolution α-tubulin image. (B) Magnification of a cell (top left) with axon filaments running across (right). (C) Magnification of part of the cell and axon filaments. (D) Cross-section of an axon bundle. Scale bar (A) 20 μm, (B) 5 μm, (C) 2 μm, (D) 0.25 μm.

**Table S1: Parameters for NN training and prediction.**

Training

| Neural network | Raw frame size | Emitter density of raw frames | Patch size | # of patches | Emitter density |
|---|---|---|---|---|---|
| DeepSTORM v1.12 | 512 x 512 | 0.028 em/µm² | 16 x 16 | 30 000 | 2 em/µm² |

| Server | GPU | # of epochs | Batch size | Learning rate | Upsamplin g factor | Percentage validation | Training time |
|---|---|---|---|---|---|---|---|
| Google Colab Pro | Tesla V100 | 100 | 256 | $1e^{-5}$ | 8 | 0.15 | 35 mins |

Prediction

| Neural network | Server | GPU | Frame size | # of frames | Prediction time |
|---|---|---|---|---|---|
| DeepSTORM v1.13 | Google ColabPro/Colab | Tesla P100/ Tesla K80 | 512 x 512 | 400 | 7 - 25 mins |
